# ESCargo: a regulatable fluorescent secretory cargo for diverse model organisms

**DOI:** 10.1101/2020.09.17.302133

**Authors:** Jason C. Casler, Allison L. Zajac, Fernando M. Valbuena, Daniela Sparvoli, Okunola Jeyifous, Aaron P. Turkewitz, Sally Horne-Badovinac, William N. Green, Benjamin S. Glick

## Abstract

Membrane traffic can be studied by imaging a cargo protein as it transits the secretory pathway. The best tools for this purpose initially block exit of the secretory cargo from the endoplasmic reticulum (ER), and then release the block to generate a cargo wave. However, previously developed regulatable secretory cargoes are often tricky to use or specific for a single model organism. To overcome these hurdles for budding yeast, we recently optimized an artificial fluorescent secretory protein that exits the ER with the aid of the Erv29 cargo receptor, which is homologous to mammalian Surf4. The fluorescent secretory protein forms aggregates in the ER lumen and can be rapidly disaggregated by addition of a ligand to generate a nearly synchronized cargo wave. Here we term this regulatable secretory protein ESCargo (<u>E</u>rv29/Surf4-dependent <u>S</u>ecretory <u>Cargo</u>) and demonstrate its utility not only in yeast cells, but also in cultured mammalian cells, *Drosophila* cells, and the ciliate *Tetrahymena thermophila*. Kinetic studies indicate that rapid transport out of the ER requires recognition by Erv29/Surf4. By choosing an appropriate ER signal sequence and expression vector, this simple technology can likely be used with many model organisms.

## Introduction

Our knowledge of the secretory pathway has progressively extended beyond morphological observations to studies of the underlying molecular machinery. Most of the key players have been identified, and they are being characterized at the biochemical and structural levels. Yet fundamental questions remain about how these components work together to drive membrane traffic. In this regard, a powerful technique is the tracking of secretory cargoes in live cells using fluorescence microscopy (Lippincott-Schwartz *et al.*, 2000). A natural or artificial secretory cargo is typically labeled with a fluorescent protein. The optimal approach is to trap the secretory cargo initially in the ER, for two reasons. First, a residence period in the ER gives the fluorescent protein portion of the cargo time to acquire a mature chromophore. Second, when the accumulated cargo is released to allow ER exit, the resulting wave of transport illuminates the stages of cargo movement through the secretory pathway (Trucco *et al.*, 2004; Boncompain and Perez, 2013).

Several regulatable secretory cargoes have been generated for mammalian cells. Tagged versions of the thermosensitive tsO45 mutant of the viral glycoprotein VSV-G accumulate in the ER at 40°C, and can be released for ER exit by dropping the temperature to 32°C (Presley *et al.*, 1997; Scales *et al.*, 1997). Similarly, procollagen accumulates in the ER at 40°C, and can be released for ER exit by reducing the temperature and adding ascorbic acid (Mironov *et al.*, 2001). These regulatable secretory cargoes are unlikely to be suitable for other model organisms. A newer approach is retention using streptavidin “hooks” (RUSH) (Boncompain *et al.*, 2012; Boncompain and Perez, 2013; Chen *et al.*, 2017). Streptavidin is fused to a resident ER protein, and it traps a secretory cargo that is tagged with the streptavidin-binding peptide (SBP). Addition of biotin to the medium inhibits the streptavidin-SBP interaction and releases the secretory cargo for ER exit. RUSH has the advantage of being compatible with a variety of natural secretory cargoes, but the initial trapping requires a low biotin concentration. In our hands, this limitation has prevented RUSH from being adapted to yeast cells (data not shown), and similar issues might arise with other model organisms.

A more versatile regulatable secretory cargo was described by Rivera *et al*. (2000). They fused GFP to four copies of the reversibly dimerizing F36M mutant of FK506-binding protein (FKBP). When this construct was targeted to the ER, dimerization of the FKBP domains created aggregates, which could be dissolved by adding a ligand that interfered with FKBP dimerization. We adapted this approach for yeast, with modifications. Improved FKBP variants with F36L and I90V mutations exhibit an increased affinity for ligands and faster disaggregation (Barrero *et al.*, 2016). When a single copy of such an FKBP variant, termed FKBP^RD^(C22V), was fused to the highly soluble tetrameric fluorescent protein DsRed-Express2 (Strack *et al.*, 2008), the result was red fluorescent aggregates that could be dissolved readily upon addition of SLF, a synthetic ligand of FKBP (Figure 1A) (Holt *et al.*, 1993; Barrero *et al.*, 2016; Casler *et al.*, 2019). When this construct was targeted to the ER lumen, fluorescent aggregates formed without causing detectable cellular stress, and addition of SLF dissolved the aggregates to produce soluble tetramers that exited the ER (Casler *et al.*, 2019).

**FIGURE 1:**
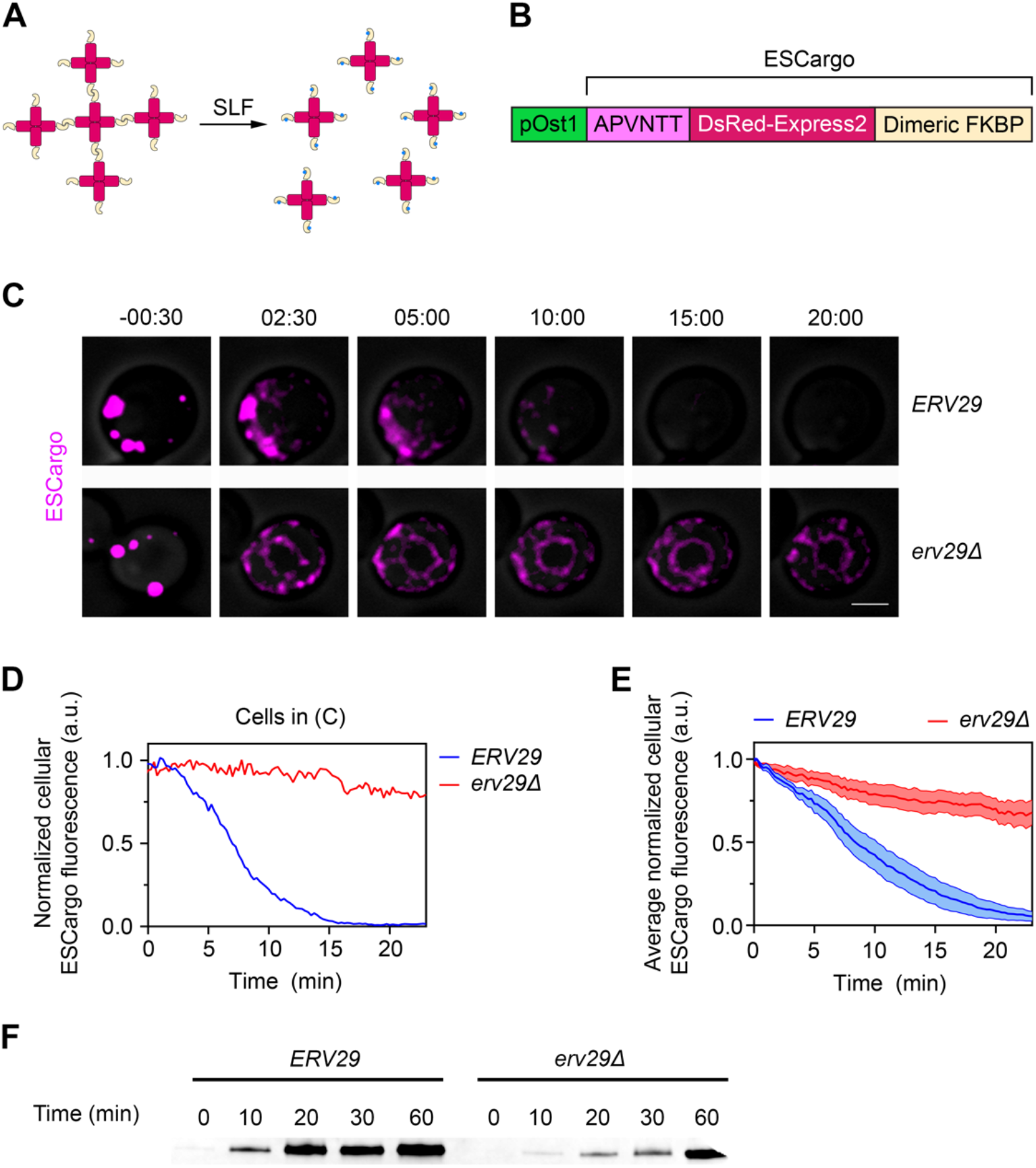
ESCargo as a tool for monitoring secretion in *Saccharomyces cerevisiae*. (A) Strategy for generating and dissolving fluorescent aggregates. DsRed-Express2 tetramers (red) fused to a dimeric variant of FKBP (gold) associate to form aggregates. Addition of the FKBP ligand SLF (blue) blocks dimerization, thereby dissolving the aggregates into soluble tetramers. (B) Functional segments of the ESCargo construct. pOst1 (green): ER signal sequence that directs cotranslational translocation in yeast. APVNTT (pink): tripeptide ER export signal followed by tripeptide *N*-linked glycosylation signal. DsRed-Express2 (red): tetrameric red fluorescent protein. Dimeric FKBP (gold): reversibly dimerizing FKBP^RD^(C22V) variant. The lengths of the segments are not to scale. (C) Secretion of ESCargo in yeast cells containing or lacking Erv29. *ERV29* wild-type and *erv29Δ* strains expressing ESCargo were grown overnight to mid-log phase and then imaged by confocal and brightfield microscopy. SLF was added at time zero to a final concentration of 100 μM. Average projected Z-stacks are taken from Supplemental Movie S1. Scale bar, 2 μm. (D) Quantification of the intracellular ESCargo fluorescence from the cells in (C). At each time point, the brightfield image was used to select the cell profile and quantify the total ESCargo fluorescence. Each fluorescence trace was normalized to the average of the three highest fluorescence signals in that trace. (E) Average intracellular ESCargo fluorescence. For each of the two strains, signals were measured from at least 8 cells from 3 movies, and the traces normalized as in (D) were averaged. Error ranges represent SEM. (F) Detecting secretion of ESCargo into the medium by immunoblotting. The strains imaged in (C) were grown overnight in rich medium, then washed and resuspended in fresh medium to the same optical density. SLF was then added to a final concentration of 100 μM to dissolve the ESCargo aggregates. After the indicated times, secreted ESCargo was precipitated from the culture medium, subjected to endoglycosidase H treatment to trim *N*-glycans, and analyzed by immunoblotting with an anti-FKBP antibody.

ER export of the DsRed-Express2-FKBP^RD^(C22V) secretory cargo was expected to occur at the relatively slow rate of bulk flow (Barlowe and Helenius, 2016). A crucial enhancement was to fuse the tripeptide APV to the N-terminus of the mature secretory cargo. This tripeptide is recognized by the Erv29 cargo receptor, and it confers rapid transport from the ER to the Golgi (Barlowe and Helenius, 2016; Yin *et al.*, 2018). The final fusion protein consisted of a cleavable ER signal sequence that drove cotranslational translocation into the ER lumen, followed by APV, followed by the tripeptide *N*-glycosylation signal NTT, followed by DsRed-Express2, followed by FKBP^RD^(C22V) (Figure 1B) (Casler and Glick, 2019). Here we designate the mature fusion protein “ESCargo” for <u>E</u>rv29-dependent <u>S</u>ecretory <u>Cargo</u>. Addition of SLF generated a wave of fluorescent ESCargo that could be tracked in maturing Golgi cisternae (Casler *et al.*, 2019). This approach was subsequently extended by adding a C-terminal vacuolar targeting peptide to ESCargo, thereby enabling a kinetic analysis of biosynthetic traffic to the yeast vacuole (Casler and Glick, 2020).

We anticipated that the original secreted version of ESCargo could be used in other cell types. In particular, mammalian cells contain an Erv29-related ER export receptor called Surf4 that recognizes the same type of tripeptide signals (Mitrovic *et al.*, 2008; Yin *et al.*, 2018). To adapt ESCargo for use in a given organism, the requirement is to append an appropriate ER signal sequence and then express this construct using a suitable vector. By comparing ESCargo with a version that lacks the tripeptide recognized by Erv29/Surf4, the kinetics can be measured for bulk flow ER export versus signal-dependent ER export (Casler *et al.*, 2019). We now document the broad utility of ESCargo as a regulatable fluorescent secretory cargo.

## Results and Discussion

### ESCargo undergoes signal-dependent ER export in yeast

ESCargo was developed to track secretion in yeast (Casler and Glick, 2019; Casler *et al.*, 2019). This protein was targeted to the ER by the Ost1 signal sequence, which drives cotranslational translocation (Willer *et al.*, 2008; Fitzgerald and Glick, 2014), thereby ensuring that fluorescent aggregates form in the ER lumen.

For the present analysis, we used a modified assay to verify that ER export of disaggregated ESCargo is accelerated by Erv29 (Casler *et al.*, 2019). Instead of measuring loss of ER-associated ESCargo by taking static images after SLF addition (Casler *et al.*, 2019), we tracked the total fluorescence signals in individual cells using 4D confocal microscopy, and then averaged the resulting traces. Although the expression vector was integrated into the genome as a single copy, there was cell-to-cell variation in the amount of aggregated ESCargo in the ER, and we reasoned that high levels of ESCargo would saturate the ER export system. Therefore, the experiment focused on cells containing moderate amounts of aggregated ESCargo. The quantification was performed in parallel using *ERV29* wild-type and *erv29Δ* strains.

Figure 1C and Supplemental Movie S1 show images of *ERV29* and *erv29Δ* cells containing ESCargo. After SLF is added, this drug must be present continuously inside the cells to prevent reaggregation. Yeast cells have pleiotropic drug transporters, so the strains carried deletions of *PDR1* and *PDR3*, which encode transcription factors that drive expression of multiple drug transporters (Schüller *et al.*, 2007; Coorey *et al.*, 2015; Barrero *et al.*, 2016). The strains also carried the *vps10-104* allele, which prevents fluorescent proteins from being diverted to the vacuole by the sortilin homolog Vps10 (Fitzgerald and Glick, 2014; Casler *et al.*, 2019). After SLF addition at time zero, the ESCargo aggregates dissolved. In the *ERV29* cell, red fluorescence was greatly diminished after 10 min and undetectable after 20 min (Figure 1C,D). By contrast, in the *erv29Δ* cell, red fluorescence persisted in the ER during the entire time course (Figure 1, C and D).

Figure 1E shows fluorescence signals averaged for multiple cells from each strain. For the *ERV29* strain, ESCargo levels began to drop soon after addition of SLF. This effect reflects rapid ER export followed by transport through the Golgi to the plasma membrane (Casler *et al.*, 2019). For the *erv29Δ* strain, the signal declined much more gradually. Presumably, ER export in this strain occurred at the slow rate of bulk flow. In support of this interpretation, an immunoblot of ESCargo in the culture medium confirmed that secretion was rapid for the *ERV29* strain but much slower for the *erv29Δ* strain (Figure 1F). Thus, ESCargo undergoes signal-dependent ER export mediated by Erv29 in a wild-type strain, or bulk flow ER export in an *erv29Δ* strain.

### ESCargo undergoes signal-dependent ER export in cultured mammalian cells

To express ESCargo in mammalian cells and target it to the ER, we subcloned the ESCargo gene cassette into a vector containing the EF-1α promoter followed by an IgH signal sequence (Strack *et al.*, 2009b). A control ESCargo variant lacked both the tripeptide ER export signal and the tripeptide *N*-glycosylation signal. That control variant, here termed ESCargo* (Figure 2A), has no known signals for ER export and was therefore expected to be a bulk flow secretory cargo. Both constructs were transfected into a Flp-In 293 T-REx cell line, which stably expressed the Golgi marker GalNAc-T2-GFP to label the juxtanuclear Golgi ribbon (Storrie *et al*., 1998).

**FIGURE 2:**
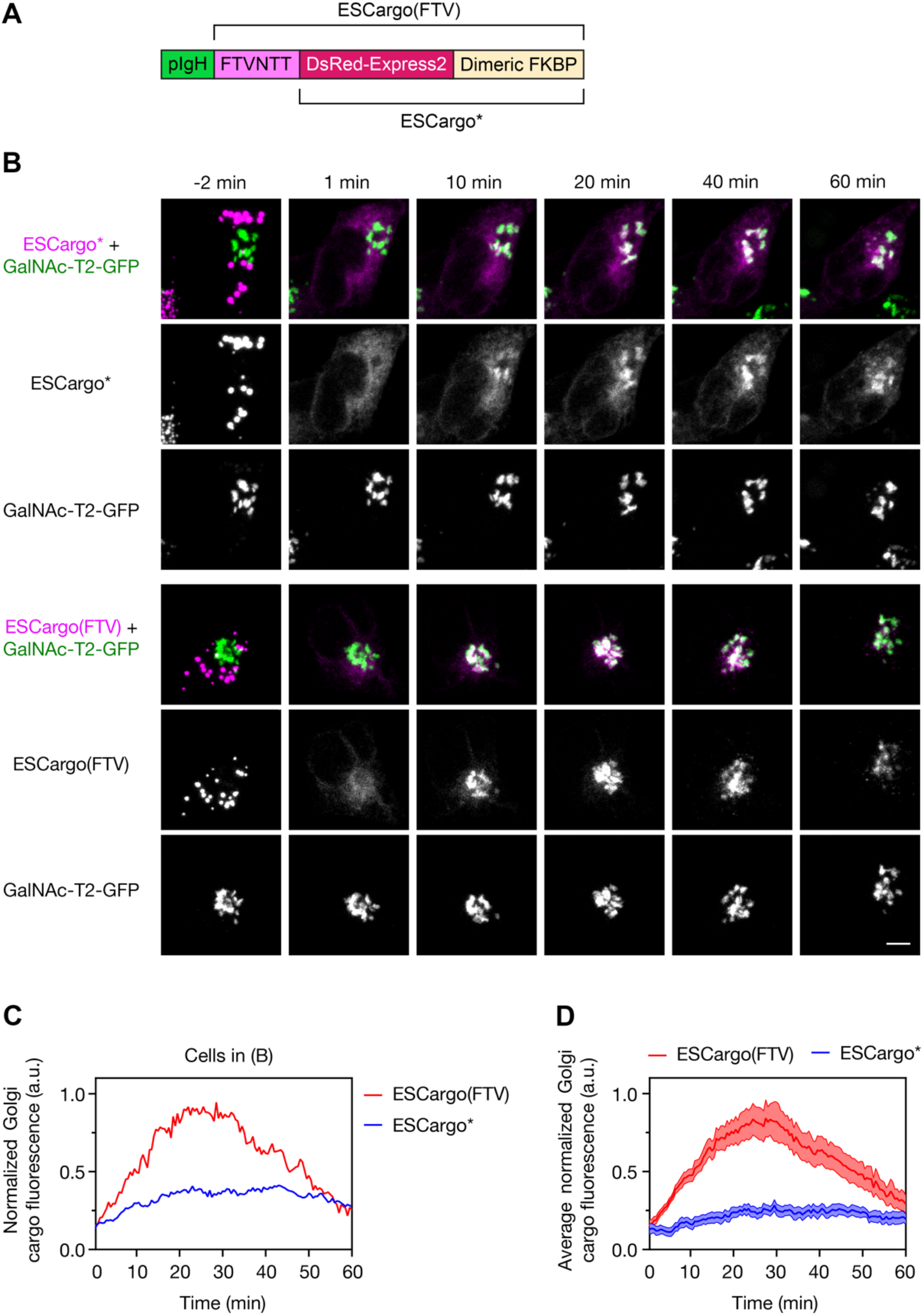
Traffic of ESCargo variants in cultured mammalian cells. (A) Functional segments of the ESCargo(FTV) and ESCargo* constructs. pIgH (green): mammalian ER signal sequence. FTVNTT (pink): tripeptide ER export signal followed by tripeptide *N*-linked glycosylation signal. For further details, see Figure 1A. (B) Comparison of the bulk flow ESCargo* variant with signal-containing ESCargo(FTV). Flp-In 293 T-REx cells stably expressing the Golgi marker GalNAc-T2-GFP were grown on confocal dishes and transfected with expression constructs for ESCargo* (top) or ESCargo(FTV) (bottom) 24-48 h before confocal imaging. Following cycloheximide treatment, SLF was added at time zero to a final concentration of 50 μM. For each cargo variant, the top row shows the merged images while the other two rows show the red and green channels. Average projected Z-stacks were taken from the first part of Supplemental Movie S2. Scale bar, 5 μm. (C) Quantification of Golgi-associated cargo fluorescence for the cells in (B). The GalNAc-T2-GFP signal was used to create masks to quantify the Golgi-associated fluorescence in the cargo channel. (D) Quantification was performed as in (C) to generate averaged time courses, using at least 7 cells from 6 movies for each variant. Error ranges represent SEM.

When the cells expressed ESCargo*, nearly all of them accumulated brightly red fluorescent round aggregates that dissolved upon SLF addition to fill the ER. A representative cell is shown in Figure 2B and Supplemental Movie S2, with the data quantified in Figure 2C. In this cell, ESCargo* fluorescence began to appear in the Golgi by 10 min after SLF addition, and reached a plateau level in the Golgi of about 20-25% of the total initial fluorescence by 20 min. The cargo signal in the Golgi remained at nearly the same level for at least 60 min. Meanwhile, substantial cargo signal remained in the ER for the entire time course. The cells had been treated with cycloheximide to suppress new protein synthesis, so the persistent ER signal was apparently due to slow ER export. These results were typical of the cells in the population, as indicated by the averaged data in Figure 2D. Thus, ESCargo* acts as expected for a bulk flow secretory cargo.

When the cells instead expressed ESCargo containing the APV tripeptide, punctate red fluorescence was observed, but for most of the cells, the fluorescence pattern showed little change upon SLF addition (data not shown). This effect highlights a limitation of ESCargo: as previously documented for yeast cells, there is a kinetic competition between aggregation in the ER and signal-dependent ER export (Casler *et al.*, 2019; Casler and Glick, 2020). In mammalian cells, ER export tends to win the race, and so much of the ESCargo accumulates in post-ER compartments. This problem was overcome by using a less potent ER export signal (Yin *et al.*, 2018). The FTV tripeptide (Figure 2A) gave satisfactory results, with nearly all of the cells exhibiting fluorescent aggregates that dissolved upon SLF addition to fill the ER (Figure 2B and Supplemental Movie S2). We therefore used the FTV variant of ESCargo, here termed ESCargo(FTV), for further experiments with mammalian cells.

After SLF addition, ESCargo(FTV) rapidly exited the ER and accumulated in the Golgi, with about 80% of the total initial fluorescence in the Golgi by 20 min after SLF addition (Figure 2 and Supplemental Movie S2). Then ESCargo(FTV) fluorescence in the Golgi progressively declined. Starting at ~25 min after SLF addition, small, mobile ESCargo(FTV)-containing structures were visible near the Golgi. These structures were presumably secretory carriers (Hirschberg *et al.*, 1998; Toomre *et al.*, 1999; Polishchuk *et al.*, 2000). At the 60-min time point, most of the ESCargo(FTV) fluorescence had left the cells, although some signal remained in the Golgi (Figure 2 and Supplemental Movie S2). The combined results indicate that ESCargo(FTV) transits the mammalian secretory pathway in a manner that involves receptor-dependent ER export.

A challenging cell type for imaging secretory cargo traffic is neurons, because the dendrites contain isolated “Golgi outposts” (reviewed in Valenzuela *et al.*, 2020). These outposts can arise by fission of tubules that extend from the somatic Golgi into the dendrites (Quassollo *et al.*, 2015) (Figure 3A and Supplemental Movie S2). We used ManII-GFP to label both the somatic Golgi and Golgi outposts in cultured rat cortical neurons (Figure 3, A and B). When ESCargo(FTV) was also expressed in those cells, spherical aggregates were present in both the soma and the dendrites (Figure 3, A and B). Within 10 min after SLF addition, ESCargo(FTV) was visible in the somatic Golgi and in all of the dendritic Golgi outposts (Figure 3, A-C and Supplemental Movie S2). Thus, ESCargo(FTV) is suitable for characterizing the secretory activity of neuronal Golgi structures.

**FIGURE 3:**
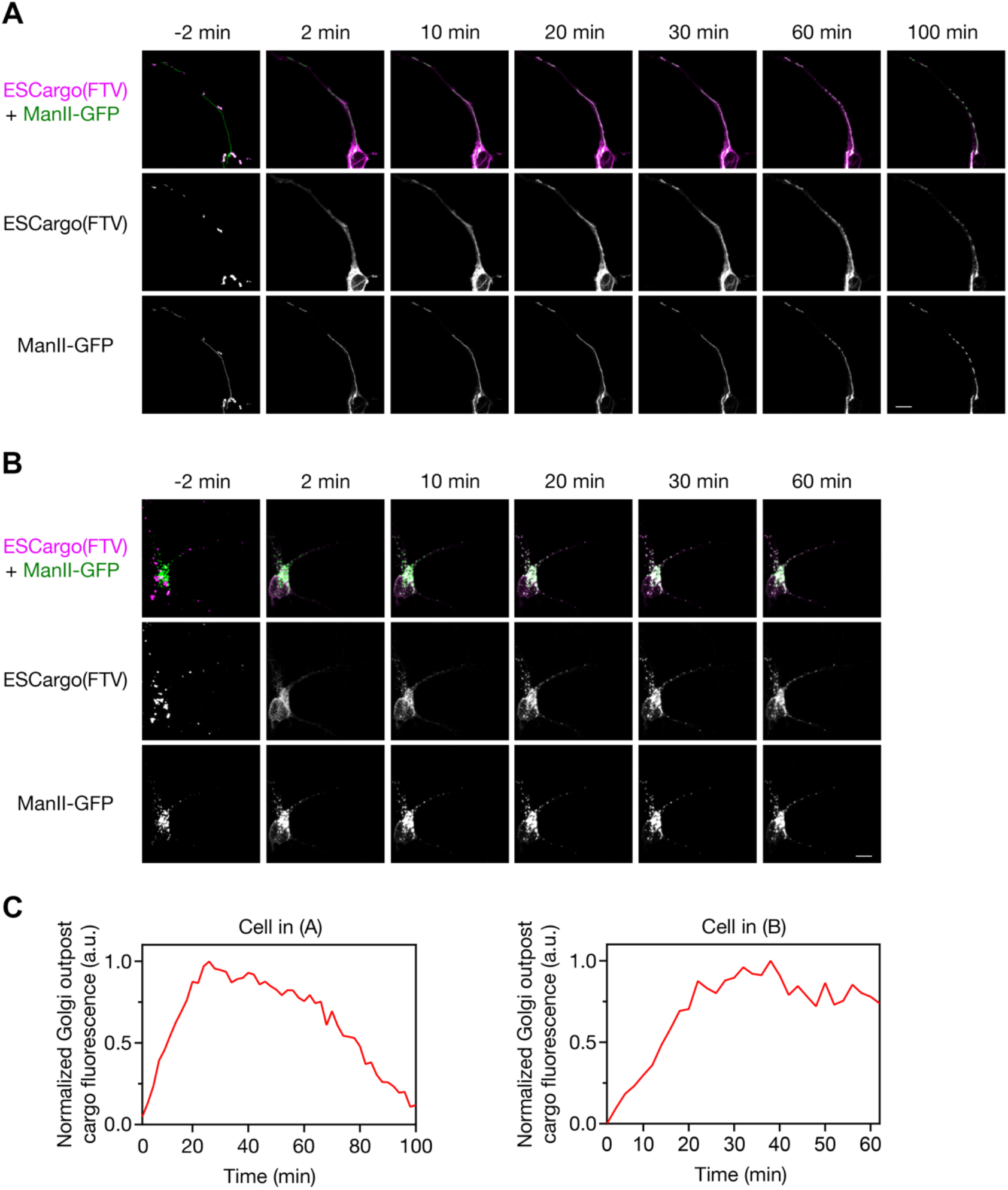
Traffic of ESCargo(FTV) in rat cortical neurons. Isolated neurons were cultured for 15 days *in vitro*, and were transfected with constructs encoding ManII-GFP and ESCargo(FTV) 48 h before imaging. SLF was added at time zero to a final concentration of 50 μM. For each of two representative cells, the top row shows the merged images while the other two rows show the red and green channels. Average projected Z-stacks were taken from the second part of Supplemental Movie S2. (A) Example of a neuron with a long tubular extension from the somatic Golgi into a dendrite. Two Golgi outposts were initially visible, and the tubule progressively fragmented to create additional Golgi outposts. Scale bar, 10 μm. (B) Example of a neuron that initially contained multiple punctate Golgi outposts. Scale bar, 10 μm. (C) Quantification of Golgi outpost-associated cargo fluorescence in the cells in (A) and (B). The ManII-GFP signal was used to create masks to quantify the Golgi outpost-associated fluorescence in the cargo channel.

### ESCargo undergoes signal-dependent ER export in *Drosophila* cells

In a procedure similar to the one described for mammalian cells, the *Drosophila* BiP signal sequence (Ohmuro-Matsuyama and Yamaji, 2018) was fused to ESCargo* and ESCargo. These two constructs were expressed in *Drosophila* S2 cells together with the Golgi marker ManII-GFP, which labeled multiple individual Golgi stacks (Zhou *et al.*, 2014).

Both ESCargo* and ESCargo formed red fluorescent round aggregates that dissolved rapidly upon addition of SLF (Figure 4A and Supplemental Movie S3). ESCargo* was visible in the ER throughout the time course, and was also persistently visible in the Golgi starting at 5-10 min. By contrast, ESCargo showed only a very transient distribution throughout the ER, and then moved completely to the Golgi within 2 min (Figure 4, A and B, and Supplemental Movie S3). Most of the ManII-GFP-labeled structures acquired ESCargo fluorescence. The remaining ManII-GFP-labeled structures were probably not Golgi stacks, because separate triple-label experiments (Figure S1A) indicated that some ManII-GFP-labeled structures were distant from the ER exit site marker Tango1 and the Golgi marker GM130 (Liu *et al.*, 2017). Indeed, when GM130 was used as the Golgi marker, all of the Golgi stacks showed cargo accumulation (Figure S1B). Starting at about 10 min after SLF addition, ESCargo began to depart from the Golgi stacks in small mobile carriers, and by 20 min, the cellular ESCargo fluorescence was almost completely gone (Figure 4, A and B, and Supplemental Movie S3). These results indicate that ESCargo exits the *Drosophila* ER in a signal-dependent manner and then rapidly traverses the secretory pathway.

**FIGURE 4:**
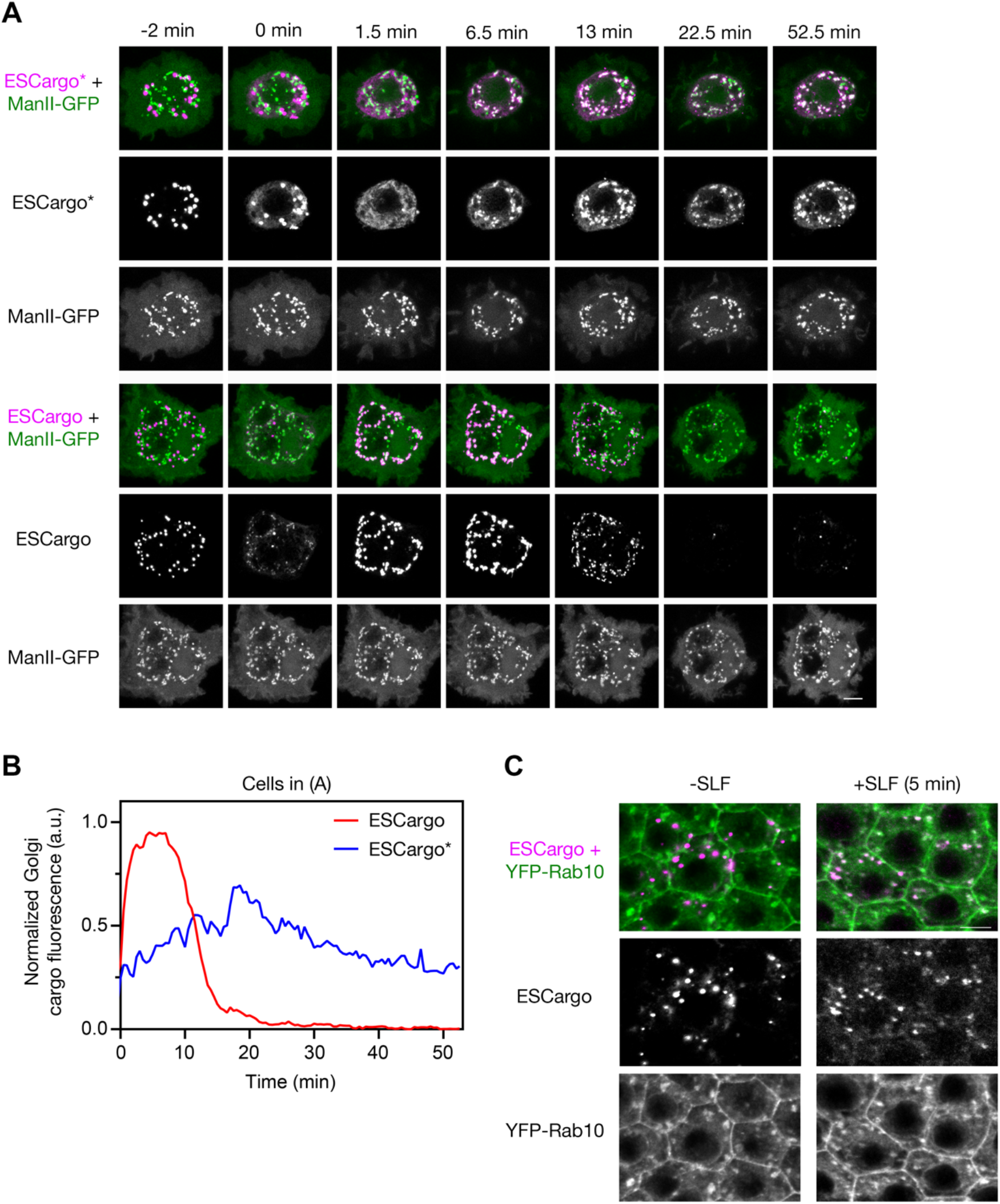
Traffic of ESCargo variants in *Drosophila melanogaster*. (A) Comparison of the bulk flow ESCargo* variant with signal-containing ESCargo in *Drosophila* S2 cells. Cells were transfected with Ubi-GAL4, pUASt-ManII-eGFP, and either pUASt-ssBiP-ESCargo* (top) or pUASt-ssBiP-ESCargo (bottom). After 3-4 days, the cells were adhered to ConA-coated confocal dishes for 30 min before confocal imaging. SLF was added at time zero to a final concentration of 50 μM. For each cargo variant, the top row shows the merged images while the other two rows show the red and green channels. Average projected Z-stacks were taken from Supplemental Movie S3. Scale bar, 5 μm. (B) Quantification of Golgi-associated cargo fluorescence for the cells in (A). The ManII-GFP signal was used to create masks to quantify the Golgi-associated fluorescence in the cargo channel. (C) Colocalization of ESCargo with the Golgi in *Drosophila* egg chamber follicular epithelial cells. Egg chambers from a *Drosophila* line (w; traffic jam-Gal4/+; UASt-ssBiP-ESCargo/UASp-YFP-Rab10) expressing ESCargo and YFP-Rab10 were fixed before and 5 min after introducing 50 μM SLF. Shown are average projections of the central four slices from confocal Z-stacks. The top row shows the merged images while the other two rows show the red and green channels. Scale bar, 5 μm.

*Drosophila* also presented an opportunity to test whether ESCargo could be used in a multicellular organism. We generated a *Drosophila* line in which the ER-targeted ESCargo construct had been inserted on chromosome 3R. Expression in follicular epithelial cells in the egg chamber resulted in large red fluorescent aggregates (Figure 4C). After incubation with SLF for 5 min, much of the red fluorescence had redistributed to areas marked by YFP-Rab10, which clusters near Golgi stacks (Figure 4C) (Lerner *et al.*, 2013). This result suggests that ESCargo behaves similarly in cultured cells and in an intact tissue.

### ESCargo* undergoes regulated secretion in *Tetrahymena thermophila*

Our analysis had been limited to opisthokonts, which are closely related from an evolutionary standpoint. We therefore turned to the evolutionarily distant ciliate *Tetrahymena thermophila*. In addition to its specialized organelles, *Tetrahymena* contains standard secretory pathway organelles including the ER and Golgi, and this model organism has been used extensively to study membrane traffic (Nusblat *et al.*, 2012).

For translocation into the *Tetrahymena* ER, we used the signal sequence of the mucocyst protein Grl1 (Chilcoat *et al.*, 1996). The cleavage site of this signal sequence has not been experimentally determined, so it was not possible to create a construct in which the APV tripeptide would be reliably located at the N-terminus of the mature protein. Instead, the Grl1 signal sequence was fused to ESCargo*. This construct was expressed under control of a cadmium-inducible promoter (Shang *et al.*, 2002).

No fluorescence was seen before cadmium induction (data not shown), but after induction, the cells contained scattered punctate fluorescent structures that were presumably ER-localized aggregates (Figure 5A). The puncta remained stable for at least an hour (data not shown). After addition of SLF, the puncta disappeared, and the cells exhibited dispersed fluorescence in an ER-like pattern throughout a 30-min time course (Figure 5A). By 30 min, some punctate fluorescence had returned (Figure 5A), perhaps due to reaggregation caused by extrusion or degradation of SLF. Within 5 min of adding SLF, secreted protein could be detected in the medium by immunoblotting (Figure 5B). A full analysis would involve experiments beyond the scope of this study, including identification of the signal sequence cleavage site, testing of putative tripeptide ER export signals, colocalization tests with organelle markers, and a time course of secretion. However, the available results suggest that ESCargo* can be secreted in *Tetrahymena*.

**FIGURE 5:**
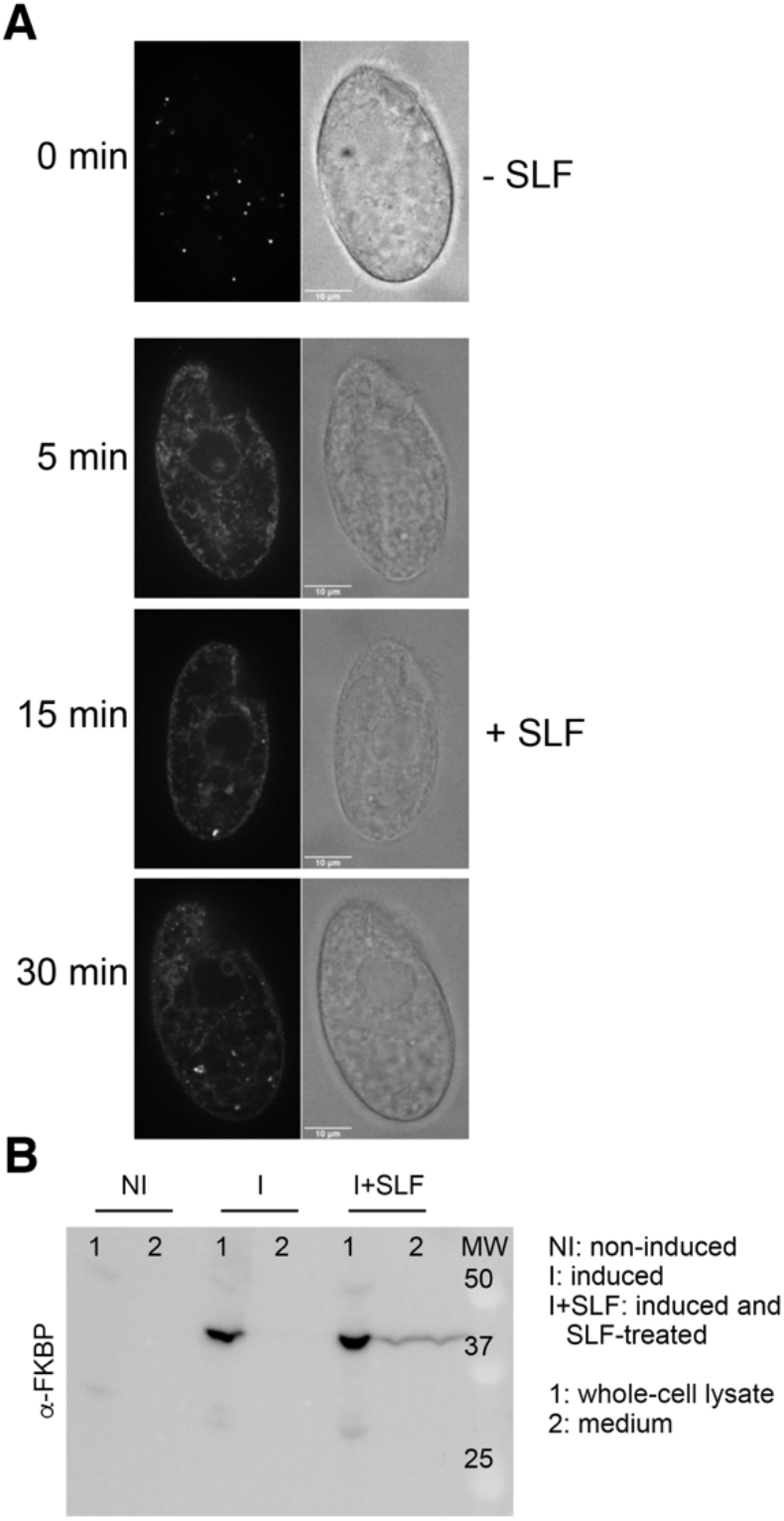
Traffic of ESCargo* in *Tetrahymena thermophila*. (A) Confocal cross sections of fixed *Tetrahymena* cells expressing ER-targeted ESCargo*, with paired differential interference contrast images. Protein expression was induced with CdCl_2_ prior to addition of 12.5 μM SLF. The top panel shows cells fixed immediately after SLF addition (0 min), and the other panels show cells fixed after treatment with SLF for 5, 15, or 30 min. The fluorescence exposure times were 100 ms for the 0 min image or 400 ms for the other images. Bright fluorescent puncta were visible initially but disappeared within 5 min after SLF addition, resulting in dispersed fluorescence in ER-like membranes that included the nuclear envelope. By 30 min, some punctate fluorescence had reappeared. Scale bars, 10 μm. (B) Immunoblot analysis of ESCargo* secretion. *Tetrahymena* cells expressing ESCargo* were treated with 12.5 μM SLF for 5 min. After centrifugation, TCA-precipitated cell pellet and cell-free culture medium samples were analyzed by SDS-PAGE and immunoblotting with anti-FBKP antibody. The cargo protein was detected in whole cell lysates for induced untreated and induced SLF-treated cells, and in the medium only for induced SLF-treated cells. No cargo protein was detected in non-induced cells.

## Conclusions

Our analysis shows that ESCargo can be used as a regulatable red fluorescent secretory cargo in model organisms. If a different fluorescent color is desired, the DsRed-Express2 portion of the cargo can be replaced with a DsRed-Express2 variant, either the orange E2-Orange (Strack *et al.*, 2009a) or the far-red E2-Crimson (Strack *et al.*, 2009b). We have seen good results with those variants (data not shown). In *S. cerevisiae, Drosophila*, and mammalian cells, SLF can be used to trigger rapid dissolution and Erv29/Surf4-dependent ER export of ESCargo. It is unknown whether evolutionarily distant organisms such as plants have Erv29/Surf4 homologs that recognize tripeptide ER export signals. However, initial results with the ciliate *Tetrahymena thermophila* suggest that in a wide range of eukaryotes, ESCargo aggregates can be generated in the ER and then dissolved with SLF to trigger secretion.

The ESCargo method has two limitations. First, when ESCargo molecules containing the APV tripeptide enter the ER, they experience a kinetic competition between aggregation and signal-dependent ER export (Casler *et al.*, 2019; Casler and Glick, 2020). As a result, a substantial fraction of the newly synthesized ESCargo molecules are either secreted or delivered to post-ER compartments. Although this phenomenon did not interfere with our studies of yeast, it did prevent efficient trapping of ESCargo aggregates in mammalian cells. We solved this problem by using the FTV tripeptide, which is a less potent ER export signal (Yin *et al.*, 2018) that allows for aggregate formation while still conferring rapid ER export of the solubilized ESCargo tetramers. Other tripeptide ER export signals might be optimal for different organisms. Second, some cells efficiently extrude SLF, thereby preventing complete and sustained dissolution of ESCargo aggregates. We overcame this hurdle for *S. cerevisiae* by blocking the expression of pleiotropic drug transporters (Barrero *et al.*, 2016; Casler and Glick, 2019), but in the budding yeast *Pichia pastoris*, we have been unable to solubilize ER-localized ESCargo aggregates with SLF (data not shown). Perhaps better results could be obtained with other FKBP ligands. In general, ESCargo is suited to cells with limited capacity to export drugs.

Our data illustrate that signal-dependent ER export can be far more efficient than bulk flow. A previous study concluded that bulk flow was surprisingly fast (Thor *et al.*, 2009). However, as pointed out by Yin *et al*. (2018), the secretory cargo used in the 2009 study contained a hemagglutinin tag (YPYDVPDYA) immediately downstream of the signal sequence, and the YPY tripeptide is recognized by Surf4 as an ER export signal, so secretion of that cargo almost certainly did not occur by bulk flow. The ESCargo* variant described here is devoid of known ER export signals and should be a good choice for measuring bulk flow secretion.

We envision that ESCargo will be useful for probing how the secretory pathway operates. The data support a cisternal maturation model for traffic through the yeast Golgi (Casler *et al.*, 2019; Kurokawa *et al.*, 2019), but the mechanism of traffic through the mammalian Golgi is ambiguous (Patterson *et al.*, 2008; Glick and Luini, 2011; Lavieu *et al.*, 2013; Rizzo *et al.*, 2013; Tie *et al.*, 2016; Dunlop *et al.*, 2017). According to a simple version of the maturation model, after secretory cargo molecules arrive at the mammalian Golgi, they should exhibit a lag period due to transit through the Golgi stack, and should then depart with linear kinetics. But according to the rapid partitioning model, after secretory cargo molecules arrive at the Golgi, they should begin to exit immediately, and should depart with exponential kinetics (Patterson *et al.*, 2008). In mammalian cells, ESCargo showed a pronounced lag period in the Golgi before appearing in secretory carriers, as predicted by the maturation model, but some ESCargo molecules persisted in the Golgi for an extended time, as predicted by the rapid partitioning model. In *Drosophila* cells, ESCargo showed a lag period before appearing in secretory carriers, and then departed from the Golgi with close to linear kinetics, fulfilling both predictions of the maturation model. These observations are preliminary and will require rigorous follow-up studies, but they offer encouragement that new tools will help to paint a unified picture of the secretory pathway.

## Materials and Methods

### Yeast growth, transformation, microscopy, and immunoblotting

The parental haploid strain was JK9-3da (*leu2-3,112 ura3-52 rme1 trp1 his4*) (Kunz *et al.*, 1993). Yeast were grown with shaking in baffled flasks at 23°C in the nonfluorescent minimal glucose dropout medium NSD (Bevis *et al.*, 2002) or in the rich glucose medium YPD supplemented with adenine and uracil. Introduction of the *vps10-104* mutation and deletion of the *PDR1* and *PDR3* genes were performed as previously described (Casler and Glick, 2019). Secretory cargo proteins were expressed using a *TRP1* integrating vector with the strong constitutive *TPI1* promoter and the *CYC1* terminator (Losev *et al.*, 2006). To ensure similar expression between strains, each clone was verified to have a single copy of the integrated plasmid by PCR using primers 5’-GTGTACTTTGCAGTTATGACG-3’ and 5’-AGTCAACCCCCTGCGATGTATATTTTCCTG-3’.

For live-cell fluorescence microscopy, yeast strains were grown in NSD (pH ~4) at 23°C. Where indicated, SLF was diluted from a 100 mM stock solution in ethanol (Cayman Chemical; 10007974) to a final concentration of 100 μM. Cells were attached to a concanavalin A (ConA)-coated coverglass-bottom dish containing NSD (Losev *et al.*, 2006), and were imaged on a Leica SP8 confocal microscope equipped with a 1.4 NA/63x oil objective using a 60 to 80 nm pixel size, a 0.25 to 0.30 μm Z-step interval, and 20-30 optical sections. Movies were deconvolved with Huygens Essential (Scientific Volume Imaging) using the classic maximum likelihood estimation algorithm, then converted to hyperstacks and average projected, then range-adjusted to maximize contrast in ImageJ (Johnson and Glick, 2019).

For immunoblotting of secreted ESCargo (Casler and Glick, 2020), each 10-mL yeast culture was grown in YPD overnight with shaking in a baffled flask to an OD_600_ of 0.8. The cells were collected by a brief spin in a microcentrifuge, washed twice with fresh YPD, and resuspended in fresh YPD to the same OD_600_. Cultures were then treated with 100 μM SLF. At each time point, a sample was processed as follows. A 1.6-mL aliquot was removed, and the cells were collected by spinning at 2500xg (5000 rpm) for 2 min in a microcentrifuge. The culture medium supernatant was transferred to a fresh microcentrifuge tube on ice, then precipitated with 4% trichloroacetic acid (TCA) on ice for 20 min. Precipitated proteins were centrifuged at maximum speed in a microcentrifuge for 15 min at 4°C. Finally, each protein pellet was resuspended in 25 μL SDS-PAGE sample buffer. Treatment with endoglycosidase H was performed as described by the manufacturer (New England Biolabs; P0702S). Briefly, Glycoprotein Denaturing Buffer was added to the protein sample, which was boiled for 5 – 10 min, followed by addition of GlycoBuffer 3 and endoglycosidase H. The incubation was at 37°C for at least 1 hr. Then 20 μL of each sample was run on a 4-20% Tris-glycine gel (Bio-Rad; 4561094). The separated proteins were transferred to a PVDF membrane (Bio-Rad; 1704156) using the Trans-Blot Turbo system (Bio-Rad). The membrane was blocked with 5% nonfat dry milk in TBST (50 mM Tris-HCl at pH 7.6, 150 mM NaCl, and 0.05% Tween 20) with shaking at room temperature for 1 h, and then incubated with 1:1000 polyclonal rabbit anti-FKBP12 antibody (Abcam; ab2918) in 5% milk/TBST with shaking overnight at 4°C. After four 5-min washes in TBST, the membrane was incubated with a 1:1000 dilution of goat anti-rabbit secondary antibody conjugated to Alexa Fluor 647 (Thermo Fisher; A21245) for 1 h at room temperature. The membrane was then washed twice with TBST and twice with 50 mM Tris-HCl at pH 7.6, 150 mM NaCl. Analysis was performed with a LI-COR Odyssey CLx imaging system.

### Mammalian cell culture, engineering, and microscopy

Flp-In 293 T-REx cells (Thermo Fisher; R78007) were maintained in 5% CO_2_ at 37°C in DMEM supplemented with 10% fetal bovine serum (GenClone; 25-550), and were regularly checked for mycoplasma contamination using a Mycoplasma Detection Kit (SouthernBiotech; 131001). Cells were transfected with 3 μg of DNA using FuGENE HD (Promega; E2311). Cells

To generate the Flp-In 293 T-REx cell line stably expressing GalNAc-T2-GFP, a synthetic gBlock (IDT) encoding a mammalian codon-optimized GalNAc-T2-GFP gene was fused to the msGFP2 gene (Valbuena *et al.*, 2020), and this construct was inserted into pcDNA5/FRT/TO (Thermo Fisher; V652020) to generate pcDNA5-GalNAc-T2-GFP. This plasmid was transfected into cells together with the Flp-recombinase expression vector pOG44 (Thermo Fisher; V600520). After two days, the medium was supplemented with 200 μg/mL hygromycin. Over the next two weeks, the medium was replaced every few days until colonies were visible on the dish. Individual clones were scraped onto a new dish and expanded. Successful integration was confirmed by visualizing GalNAc-T2-GFP fluorescence in all cells, by newly acquired sensitivity to Zeocin (400 μg/mL), and by resistance to hygromycin (200 μg/mL).

For live-cell imaging of Flp-In 293 T-Rex cells, cultures were grown on coverglass-bottom dishes coated with polyethyleneimine (PEI) (Vancha *et al.*, 2004). A dish was incubated with 250 μL of ~25 μg/mL PEI (Sigma; 181978) in 150 mM NaCl for 10 min, and then the dish was washed and allowed to dry completely before adding cells. Prior to imaging, cells were treated with 100 μg/mL cycloheximide from a 100 mg/mL stock in DMSO for 15-30 min, and the medium was buffered with 15 mM Na^+^-HEPES, pH 7.4. Imaging was performed at 37°C on a Leica SP5 confocal microscope equipped with an incubator stage and a 1.4 NA/63x oil objective, using a 100 nm pixel size, a 0.5 μm Z-step interval, and ~30-40 optical sections. Z-stacks were average projected to make movies.

For live-cell imaging of neurons, primary cultures of rat cortical neurons were prepared as described (Govind *et al.*, 2012) using Neurobasal Medium (Thermo Fisher; 21103049), 2% (v/v) B27 Supplement (Thermo Fisher; 17504044), and 2 mM L-glutamine (Thermo Fisher; 25030024). Dissociated cortical cells from E18 Sprague Dawley rat pups were plated in 35 mm glass bottom dishes (MatTek; P35G-1.5-14-C) coated with poly-D-lysine (Sigma; A-003-E) at a density of 0.25×10^6^ cells/mL. Neuronal cultures were transfected at 15 days *in vitro* with 1 μg of cDNA encoding a GFP-tagged version of the Man2A1 alpha-mannosidase II (ManII-GFP) and 1 μg of plasmid encoding ER-targeted ESCargo(FTV) per dish using 2 μL of Lipofectamine 2000 reagent (Thermo Fisher; 11668019). Neurons were incubated for 48 h post-transfection and then imaged using a Leica SP5 confocal microscope as described above. Immediately prior to imaging, the cell medium was replaced with 2.5 mL of Hibernate E (BrainBits).

ImageJ was used to quantify the Golgi-associated secretory cargo signals in the movies. The first step was to measure background cellular fluorescence from a control movie of non-transfected cells. Then for each time point, the signal from the GFP-tagged Golgi marker was used to create a mask, and the background-subtracted cargo signal within the mask was measured. For Flp-In 293 T-REx cells, these values were normalized to the initial cargo fluorescence, which was measured by quantifying the total background-subtracted cellular fluorescence at 1 min after SLF addition. For rat cortical neurons, these values were normalized to the maximal cargo signal measured in the Golgi outposts.

### *Drosophila* cell culture, engineering, and microscopy

S2-DGRC cells were obtained from the *Drosophila* Genomic Resource Center (cell line #6). Cells were grown in Insect-XPRESS™ Protein-free Insect Cell Medium (Lonza; 12-730Q) supplemented with antibiotic-antimycotic (Thermo Fisher; 15240062) in a 25°C incubator. The DNA constructs used were Ubi-GAL4 (a gift of Richard Fehon), pUASt-ManII-eGFP (a gift of Bing Ye), and either pUASt-ssBiP-ESCargo* or pUASt-ssBiP-ESCargo. Transfection was performed using the dimethyldioactadecylammonium (DDAB) method (Han, 1996). DDAB and cell culture medium were mixed in a 1:2 ratio and incubated at room temperature for 10 min. To transfect a well in a 6-well plate, 1 μg of DNA was mixed with 144 μL of the DDAB/medium solution and incubated for 15 min at room temperature, and this mixture was added to 2.6×10^6^ cells in 2 mL of medium. The transfected cells were allowed to grow for 3-4 days before imaging.

For immunofluorescence microscopy, coverslips were incubated in 100 μg/mL ConA for 1 h, washed 3x in deionized water, and dried before use. Transfected cells were detached by pipetting, and 250 μL of a cell suspension was added to a fresh 6-well plate containing a ConA-coated coverslip and 1.75 mL of media. Cells were allowed to spread for 1 h, and were then fixed in 4% formaldehyde diluted in phosphate buffered saline (PBS) for 10 min at room temperature. The fixed cells were washed, and permeabilized with PBS containing 0.1% Triton X-100 (PBST) 3x for 5 min each. Anti-GM130 antibody (Abcam; ab30637) and anti-Tango1 antibody (Lerner *et al.*, 2013) were diluted 1:500 in PBST. A 100-μL drop of the antibody mixture was placed on a piece of parafilm in a humidified chamber, and the cell-coated coverslip was inverted onto this drop and incubated for 1 h at room temperature. The coverslip was washed 3x for 5 min each in PBST. Secondary antibodies (Thermo Fisher; A-27039 and A-2145) diluted 1:500 in PBST were added for 1 h at room temperature, and the samples were washed 3x for 5 min each in PBST. Finally, the coverslip was mounted in ProLong™ Gold (Thermo Fisher; P36930) and sealed with nail polish. Imaging was performed on a Zeiss LSM 800 laser scanning confocal microscope with Zen blue software and a 63x Plan Apo 1.4 NA oil objective.

For live imaging of S2 cells, transfected cells were detached by pipetting, and 250 μL of cells and 750 μL of fresh media were added to a 35 mm glass bottom dish coated with ConA using the procedure described above for coverslips. Cells were allowed to attach for 30 min, and were then imaged at room temperature on a Zeiss LSM 880 laser scanning microscope with Zen black software and a 63x Plan Apo 1.4 NA oil objective. Every 30 s for 1 h, a Z-stack of ~2-5 μm, depending on cell height, was captured using steps of 0.37 μm. After the first 2 min of imaging, 1 mL of medium containing 100 μM SLF was added, yielding a final SLF concentration of 50 μM. The Z-stacks were average projected and quantified as described above for cultured mammalian cells.

Transgenic flies containing the pUASt-ssBiP-ESCargo plasmid were generated using PhiC31-mediated recombination. Insertions were generated on chromosomes 2L(attP40) and 3R(VK20) by GenetiVision. Both lines expressed well, and the VK20 insertion was used for all experiments in this paper. The egg chamber follicle cell driver traffic jam-Gal4 from the *Drosophila* Genetic Resource Center in Kyoto (DGRC #104055) was used to express UASt-ssBiP-ESCargo as well as UASp-YFP-Rab10 from the Bloomington *Drosophila* Stock Center (BDSC #9789).

To examine ESCargo traffic in egg chambers, transgenic flies were reared on cornmeal molasses agar at 25°C using standard techniques. 1-to 2-day old females were aged on yeast with males for 2 days at 25°C. Ovary dissection was performed as described (Cetera *et al.*, 2016). Ovaries were removed from yeasted females in dissection/live cell imaging medium (Schneider’s *Drosophila* medium containing 15% FBS and 200 μg/mL insulin). Ovariole strands were mechanically removed from muscle with forceps. Egg chambers older than stage 9 were cut away from the ovariole strand using a 27-gauge needle. To trigger a wave of ESCargo traffic, egg chambers were transferred to a 1.5 mL tube, and were incubated in fresh imaging medium for 5 min at room temperature with 50 μM SLF, or without SLF as a negative control. Egg chambers were fixed in 4% formaldehyde in PBST for 15 min at room temperature, washed 3x for 10 min each in PBST, and mounted in ~35 μL ProLong™ Gold on a slide with a 22×50mm coverslip. The coverslip was then sealed with nail polish. Imaging was performed on a Zeiss LSM 800 laser scanning confocal microscope with Zen blue software and a 63x Plan Apo 1.4 NA oil objective.

### *Tetrahymena* cell culture, engineering, and microscopy

*Tetrahymena thermophila* were grown overnight in SPP (2% proteose peptone, 0.1% yeast extract, 0.2% dextrose, 0.003% ferric-EDTA) supplemented with 250 μg/mL penicillin G, 250 μg/mL streptomycin sulfate, and 0.25 μg/mL amphotericin B fungizone, to medium density (0.6-2.0 x 10^5^ cells/mL). For biolistic transformation, growing cells were subsequently starved in 10 mM Tris buffer, pH 7.4, for 18-20 h. Fed and starved cells were kept at 30°C with agitation at 99 rpm, unless otherwise indicated. Culture densities were measured using a Z1 Coulter Counter (Beckman Coulter).

After biolistic transformation, transformants were selected as previously described (Kaur *et al.*, 2017; Sparvoli *et al.*, 2018). Transformants were serially transferred 6x per week in increasing concentrations of blasticidin and decreasing concentrations of CdCl_2_ (up to 90 μg/mL of blasticidin and 0.1 μg/mL CdCl_2_) for at least 3 weeks before further testing.

For fluorescence microscopy, cells were grown overnight in 20 mL SPP at 30°C with agitation at 99 rpm to ~2.0×10^5^ cells/mL. Transgene expression was induced with 0.5 μg/mL CdCl_2_ for 90 min. Cells were then washed once with 20 mL SPP and resuspended in 20 mL of medium without CdCl_2_, prior to treatment with SLF. Incubation with 12.5 μM SLF was for 5, 10, or 30 min in a 24-well plate (2 mL/well) at 30°C without agitation. Cells were then fixed by addition of ice-cold 4% paraformaldehyde, and incubated for 30 min at room temperature. Controls included CdCl_2_-induced cells fixed before the treatment with SLF (time zero) or kept for 60 min in SPP without SLF. Non-induced cells were grown, fixed, and imaged in parallel. Fixed cells were washed three times with PBS, mounted in 30% glycerol in PBS containing 0.1 mM Trolox (Sigma; 238813) to inhibit bleaching, and imaged at room temperature on a Marianas Yokogawa-type spinning disk inverted confocal microscope using a 100x 1.45 NA oil objective with SlideBook 6 software (Intelligent Imaging Innovations). Z-stack images were denoised and adjusted for brightness and contrast using Fiji software (Schindelin *et al.*, 2012), and individual optical sections were chosen for display.

To detect secreted proteins by immunoblotting, cells were grown overnight in 20 mL SPP to 0.6-0.8×10^5^ cells/ml. 1.0×10^5^ cells were transferred into 2 mL SPP for each experimental condition: noninduced, CdCl_2_-induced, and CdCl_2_-induced and SLF-treated. Protein expression was induced with 0.5 μg/mL CdCl_2_ for 90 min at 30°C. CdCl_2_-induced cells were washed once with SPP and resuspended in 1 mL SPP without or with 12.5 μM SLF for 5 min at room temperature. Non-induced cells were washed and resuspended in SPP for 5 min without SLF. Cells were pelleted by centrifugation, and ~500 μL of cell-free supernatant were precipitated by adding one-tenth volumes of 2% deoxycholate and 100% TCA. In parallel, the corresponding pellet fractions were precipitated with 10% TCA. TCA-insoluble pellets were suspended in 100°C LDS (lithium dodecyl sulfate) sample buffer containing 40 mM DTT, and analyzed by SDS-PAGE and immunoblotting as previously described (Sparvoli *et al.*, 2018). A rabbit monoclonal anti-FBKP antibody (Abcam; ab2918) was diluted 1:1000 in blocking solution. Proteins were visualized with 1:20,000 anti-rabbit IgG (whole molecule)-peroxidase secondary antibody (Sigma; A0545, lot 022M4811) and SuperSignal West Femto Maximum Sensitivity Substrate (Thermo Fisher; 34095).

### Plasmids

All newly generated plasmids used in this study are documented in the online supplemental ZIP file, in the form of a text document plus annotated map/sequence files that can be opened with SnapGene Viewer (Insightful Science; www.snapgene.com). All relevant inserts in the plasmids were verified by sequencing. Newly generated plasmids have been submitted to Addgene.

## Supporting information

Folder of DNA Constructs

## Acknowledgments

This work was supported by NIH grant R01 GM104010 to BSG, by NIH grant R01 GM105783 to APT, by NIH grant R01 GM136961 and American Cancer Society grant RSG-14-176 to SHB, and by NIH grant R01 DA044760 to WNG. JCC was supported by NIH training grant T32 GM007183. AZ was supported by American Heart Association fellowship 16POST2726018 and American Cancer Society fellowship 132123-PF-18-025-01-CSM. Thanks for assistance with fluorescence microscopy to Vytas Bindokas and Christine Labno at the Integrated Microscopy Core Facility, which is supported by the NIH-funded Cancer Center Support Grant P30 CA014599. The pUASt-ManII-eGFP plasmid was a gift from Bing Ye, and the Ubi-Gal4 plasmid was a gift from Rick Fehon.

## Author Contributions

JCC helped to design all of the experiments, created all of the secretory cargo plasmids, performed the experiments with *Saccharomyces* cells and cultured mammalian cells, created the figures and movies, and wrote the initial draft of the manuscript. AZ helped to design and performed all of the *Drosophila* experiments. FMV created the mammalian ManII-GFP construct, and performed the experiments with cultured neurons after optimizing the conditions. DS helped to design and performed all of the *Tetrahymena* experiments. OJ dissected rat brains, cultured the cortical neurons, and provided technical advice. APT supervised the *Tetrahymena* experiments. SHB supervised the *Drosophila* experiments. WNG supervised the experiments with cultured neurons. BSG helped to design all of the experiments, supervised the project, and revised the manuscript and figures.

## Abbreviations

ESCargo: Erv29/Surf4-dependent Secretory Cargo
FKBP: FK506-binding protein
GalNAc-T2: *N*-acetylgalactosaminyltransferase 2
ManII: mannosidase II
SLF: synthetic ligand of FKBP

---

**FIGURE S1:**
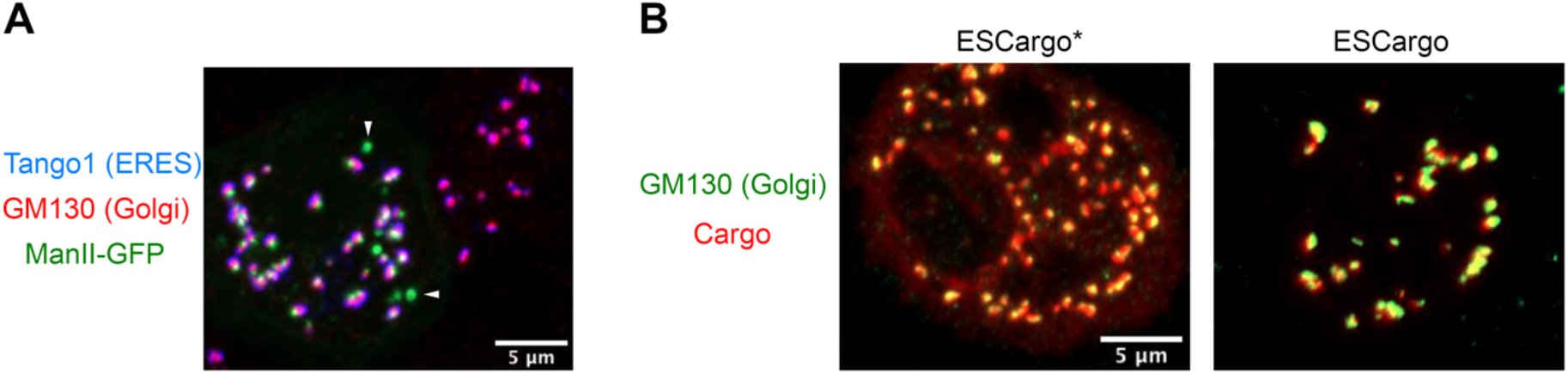
Golgi labeling in *Drosophila* S2 cells. (A) Incomplete colocalization of ManII-GFP puncta with Golgi and ER exit site (ERES) compartments. S2 cells expressing ManII-GFP were fixed and immunostained for GM130 (Golgi marker) and Tango1 (ERES marker). A subset of the ManII-GFP puncta did not have associated GM130 or Tango1 signal. Two examples are marked with arrowheads. Scale bar, 5 μm. (B) Accumulation of secretory cargo in all Golgi stacks. S2 cells expressing ESCargo* or ESCargo were treated with 50 μM SLF for 5 min, fixed, and immunostained for GM130 (Golgi marker). All of the GM130 puncta had cargo signal. Scale bar, 5 μm.

**Movie S1:** ESCargo in *Saccharomyces cerevisiae*. Shown are representative cells of a wild-type (WT) strain containing Erv29 and of an *erv29Δ*. strain. Confocal Z-stacks were captured every 15 s for 24.5 min. SLF was added at time zero to a final concentration of 100 μM. The movie frames are average projected fluorescence signals merged with brightfield images of the cells. Images from this movie are shown in Figure 1C. Scale bar, 2 μm.

**Movie S2:** ESCargo* and ESCargo(FTV) in cultured mammalian cells, and ESCargo(FTV) in rat cortical neurons. In the first part of the movie, Flp-In 293 T-REx cells stably expressing the Golgi marker GalNAc-T2-GFP were grown on confocal dishes and transfected with expression constructs for ESCargo* (left) or ESCargo(FTV) (right) 24-48 h before imaging. Following cycloheximide treatment, SLF was added at time zero to a final concentration of 50 μM. Confocal Z-stacks were taken every 30 s for 62 min. Average projections are shown for representative cells. The top panel is a merge of the two fluorescence channels. Images from this movie are shown in Fighure 2B. Scale bar, 5 μm. In the second part of the movie, rat cortical neurons were transfected to express the Golgi marker ManII-GFP together with ESCargo(FTV) 48 h before imaging. The left and right panels show separate movies of two representative cells. Confocal Z-stacks were taken 2 min prior to SLF addition, and then every 2 min after SLF addition for 100 min (left panel) or 60 min (right panel). Images from this movie are shown in Figure 3, A and B. Scale bar, 10 μm.

**Movie S3:** ESCargo* and ESCargo in *Drosophila* S2 cells. Cells were transfected with Ubi-GAL4, pUASt-ManII-eGFP, and either pUASt-ssBiP-ESCargo* (left) or pUASt-ssBiP-ESCargo (right). After 3-4 days, the cells were adhered to ConA-coated confocal dishes for 30 min before confocal imaging. SLF was added at time zero to a final concentration of 50 μM. Confocal Z-stacks were taken every 30 s for 54.5 min. Average projections are shown for representative cells. The top panel is a merge of the two fluorescence channels. Images from this movie are shown in Figure 4A. Scale bar, 5 μm.

