## Supplementary material for "ESCargo: a regulatable fluorescent secretory cargo for diverse model organisms": Folder of DNA Constructs: Table of DNA Constructs.rtf

Vector Name	Organism	Addgene Number	
YIplac204-pOst1-ESCargo	S. cerevisiae	115420	
YIplac204-pOst1-ESCargo*	S. cerevisiae	115427	
YIplac128-pOst1-ESCargo	S. cerevisiae	115421	
YIplac204-pOst1-ESCargo(Crimson)	S. cerevisiae	140154	
YIplac128-pOst1-ESCargo(Crimson)	S. cerevisiae	140156	
YIplac204-pOst1-ESCargo(Orange)	S. cerevisiae	140157	
YIplac128-pOst1-ESCargo(Orange)	S. cerevisiae	140158	
pEF-pIgH-ESCargo*	Mammals	140159	
pEF-pIgH-ESCargo(FTV)	Mammals	140160	
pEF-pIgH-ESCargo(FTV,Orange)	Mammals	140161	
pEF-pIgH-ESCargo(FTV,Crimson)	Mammals	140162	
pcDNA-GalNAc-T2-GFP-ESCargo(FTV)	Mammals	140163	
pEF-GalNAc-T2-GFP	Mammals	140164	
pEF-ManII-GFP	Mammals	XXXXXX	
pUASt-ssBiP-ESCargo* 	D. melanogaster	140165	
pUASt-ssBiP-ESCargo	D. melanogaster	140166	
NCVB-ssGrl1-ESCargo*	T. thermophila	140167	
